# TiDHy: Timescale Demixing via Hypernetworks to learn simultaneous dynamics from mixed observations

**DOI:** 10.1101/2025.01.28.635316

**Authors:** Elliott T. T. Abe, Bingni W. Brunton

## Abstract

Neural activity and behavior arise from multiple concurrent time-varying systems, including neuromodulation, neural state, and history; however, most current approaches model these data as one set of dynamics with a single timescale. Here we develop **Ti**mescale **D**emixing via **Hy**pernetworks (TiDHy) as a new computational method to model spatiotemporal data, decomposing them into multiple simultaneous latent dynamical systems that may span orders-of-magnitude different timescales. Specifically, we train a hyper-network to dynamically reweigh linear combinations of latent dynamics. This approach enables accurate data reconstruction, converges to true latent dynamics, and captures multiple timescales of variation. We first demonstrate that TiDHy can demix dynamics and timescales from synthetic data comprising multiple independent switching linear dynamical systems, even when the observations are mixed. Next, with a simulated locomotion behavior dataset, we show that TiDHy accurately captures both the fast dynamics of movement kinematics and the slow dynamics of changing terrains. Finally, in an open-source multi-animal social behavior dataset, we show that the keypoint trajectory dynamics extracted with TiDHy can be used to accurately identify social behaviors of multiple mice. Taken together, TiDHy is a powerful new algorithm for demixing simultaneous latent dynamical systems with applications to diverse computational domains.

## 1 Introduction

Over milliseconds to a lifetime, animals perceive, process, learn, respond to, and act on their environment. The computational underpinnings of these natural behaviors involve interactions among nonlinear computations across the brain and body. Therefore, data-driven methods to model natural behavior must also account for multiple simultaneous dynamical systems spanning orders-of-magnitude different scales. This computational challenge is exacerbated by the fact that measurements of animal behavior (e.g., kinematic trajectories of body keypoints) and neural activity (e.g., single-cell or population recordings of a small subset of the nervous system) are always incomplete descriptions of the true underlying systems. We refer to these true, unobserved dynamical systems as “latent” dynamics. For example, video data of mouse foraging behaviors are often processed into tracked kinematic trajectories of body parts for behavior segmentation. However, these movement trajectories reflect the synthesis of multiple overlapping, partially observed latent systems that are complex and ultimately arise from neural activity, including sensorimotor control of the musculoskeletal system, neuromodulation of arousal, social context, satiety, and behavioral history. All of these multi-scale processes simultaneously contribute to variations in behavior, yet most current methods to model these data as having dynamics over a narrow range of timescales.

From a computational perspective, we identify four primary challenges to “demix” latent systems and their contributions to experimental observations. First, latent systems cannot be measured directly, and data comprise partial or mixed observables. Second, observations may come from multiple interacting systems. Third, dynamics may be nonstationary. And fourth, the timescales of dynamics may span orders of magnitude. These challenges are not unique to modeling data from neuroscience and behavior; indeed, considerable interdisciplinary efforts in adjacent fields have been made to learn multiple latent dynamical systems from data. However, these solutions are generally tailored to the application area, tackle a subset of these challenges, and only partially address the demixing problem. We briefly review some of these modeling approaches below.

Linear approaches to the spatiotemporal data demixing problem generally pose it as a matrix decomposition, with or without the assumption that the data are generated by a dynamical process. For example, common techniques such as Gaussian mixture models [1] and independent component analysis (ICA, [2–4]) do not assume a dynamical model. These methods have been popularly used to demix and localize audio signals from different sources, including to separate multiple speakers, musical instruments, or speech from background noise. Methods like dynamic mode decomposition (DMD, [5, 6]) do assume a dynamical model and use matrix decomposition to separate spatial and temporal components to explain the data. However, a central assumption is that the latent systems are stationary in time.

Although linear methods have been successful in describing simplified systems or small windows of data, biological systems are inherently nonlinear. A popular approach is to relax the linearity assumption by allowing the solution to be non-stationary and by learning nonlinear transformations of the data. In particular, state-space models like autoregressive hidden Markov models (ARHMMs, [7–9]) and switching linear dynamical systems (SLDSs, [10]) describe dynamics with multiple discrete states. These models assume that only a single dynamical system is active at one time and has a single timescale, and they successfully reconstruct nonstationary data by also modeling the transitions among several states. To learn nonlinear dynamics explicitly, approaches like sparse identification of nonlinear dynamics (SINDy, [11–13]) have been used to infer coefficients of the nonlinear dynamic equations. Although these methods can successfully learn nonlinearities, they are most successfully applied to learn a single stationary nonlinear system. Another approach is sparse dictionary learning, which can learn combinations of simultaneous linear dynamics [14–16]. Even so, capturing multiple timescales remains a challenge, and complex nonlinear implementations often sacrifice interpretability. Among these, our proposed approach is mostly closely related to sparse dictionary learning by including a hierarchical generative model.

An alternative to explicitly learning the functional form of one or several dynamical systems is to take advantage of neural networks as universal function approximators [17]. In particular, recurrent neural networks (RNNs) have become a common model to recapitulate the dynamics of behavior [18–20] and neural activity [19, 21]. A related approach uses neural ordinary differential equation networks (Neural ODEs, [22, 23]), which have also been used to numerically solve complex differential equations of dynamical systems. Neural ODEs have been a powerful tool in applied mathematics to learn systems that can be expressed analytically; however, they have not yet been able to generalize to learn multiple latent systems. Further, neural networks can faithfully reconstruct complex nonlinear observations [24–26], but they are difficult to interpret and generalize because they act as black-box models between inputs and outputs. In addition, even as high-fidelity data reconstruction can be achieved, the learned latent variables may have little resemblance to the true dynamics of the system.

In this paper, we introduce **Ti**mescale **D**emixing via **Hy**pernetworks (TiDHy, pronounced “tidy,” as in, neatly in order) as a method to learn multiple simultaneously occurring latent dynamical systems spanning orders-of-magnitude timescales from partial or mixed observations. Our approach is to model latent dynamics as a hierarchical generative model that can dynamically reweigh combinations of linear systems from timestep to timestep. We parameterize this dynamic reweighing as a hypernetwork [26, 27], which is a compact and interpretable way to estimate the relative contributions of simultaneous dynamics. In other words, TiDHy learns how the observed data can be explained by multiple, approximately orthogonal linear systems weighted by their relative contributions. We demonstrate TiDHy’s performance to demix several datasets, as evaluated not only on reconstruction error but also on ability to identify the true latent dynamics. First, in a synthetic dataset, we show that TiDHy accurately demixes multiple independent switching linear dynamical systems (SLDSs) from partially superimposed observations. Next, using a simulated walking behavior dataset, we simultaneously capture the fast timescale of joint kinematics and the slower timescale of locomotion on changing terrain. Finally, we demonstrate that TiDHy correctly classifies natural social behavior in an open-source multianimal dataset. We suggest that TiDHy is a powerful new algorithm for disentangling many factors needed to explain natural behavior, and it can be readily applied to demix latent dynamical systems in diverse application domains.

## 2 Results

Here we establish the mathematical formulation of TiDHy as a spatiotemporal data demixing problem in Section 2.1 and introduce an algorithmic approach to learn this demixing as parameterized by a hypernetwork in Section 2.2. To build intuition, we demonstrate the interpretation of the demixed system on a series of increasingly challenging synthetic datasets for which the ground truth is known in Section 2.3. Next in Sections 2.4 and 2.5, we apply TiDHy on simulated and real behavior data that are characterized by multiple interacting dynamical systems that span multiple timescales.

### 2.1 Formulation of TiDHy

The most common framework for understanding spatiotemporal behavior and neural activity is to model recorded data as linear dynamical systems (LDS). In these methods, the goal is to learn a set of dynamics over latent states **X** that explain the experimental data **Z** at time *t*:

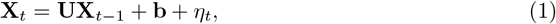

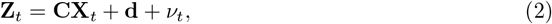

where **U** is the latent state dynamics matrix, **C** is the observation model, **b** and **d** are bias terms, and *η* and *ν* are Gaussian noise random variables.

Switching linear dynamical systems (SLDSs) are an extension of LDSs that allows for multiple sets of linear dynamics. Instead of a single **U** matrix, SLDSs model a set of **U**^(*k*)^ matrices that can probabilistically switch at every time point, so that different dynamics are active at different times. However, the SLDS model allows only a single set of dynamics to be active at each time step, so it cannot capture dynamics that require multiple simultaneous systems. To address this problem, our approach is to model the dynamics matrix as a linear combination of *K* dynamics matrices 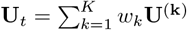 to capture simultaneous dynamical systems.

TiDHy assumes observed data **Z**_*t*_ arise from a hierarchical generative model formulated as multiple non-stationary latent dynamical systems (Fig. 1a, Supplmental Fig. A1). We model the two latent states: a higher-order latent state 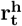, which modulates a lower-level latent dynamics **r**_**t**_ through a nonlinear transformation ℋ, parameterized as a hypernetwork [27] (Fig. 1b). In general, a hypernetwork is a network that outputs the weights of another network. By taking the higher order latent variable 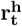 as input and outputting vector weights 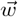 at every time point, the hypernetwork *ℋ* is thus able to modulate the transition matrices **U** of the lower-level latent variable **r**_**t**_, tracking non-stationary dynamics at different timescales. Specifically, the temporal dynamics of the latent states **r**_**t**_ are modeled as *K* learnable transition matrices 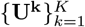, which are linearly combined according to the weight vector 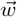. The linearly combined transition matrix **U** is then used to step the dynamics of *r* from *t* to *t* + 1. Finally, the spatiotemporal observations **Z**_**t**_ are governed by a linear observation matrix Φ.

**Fig. 1.**
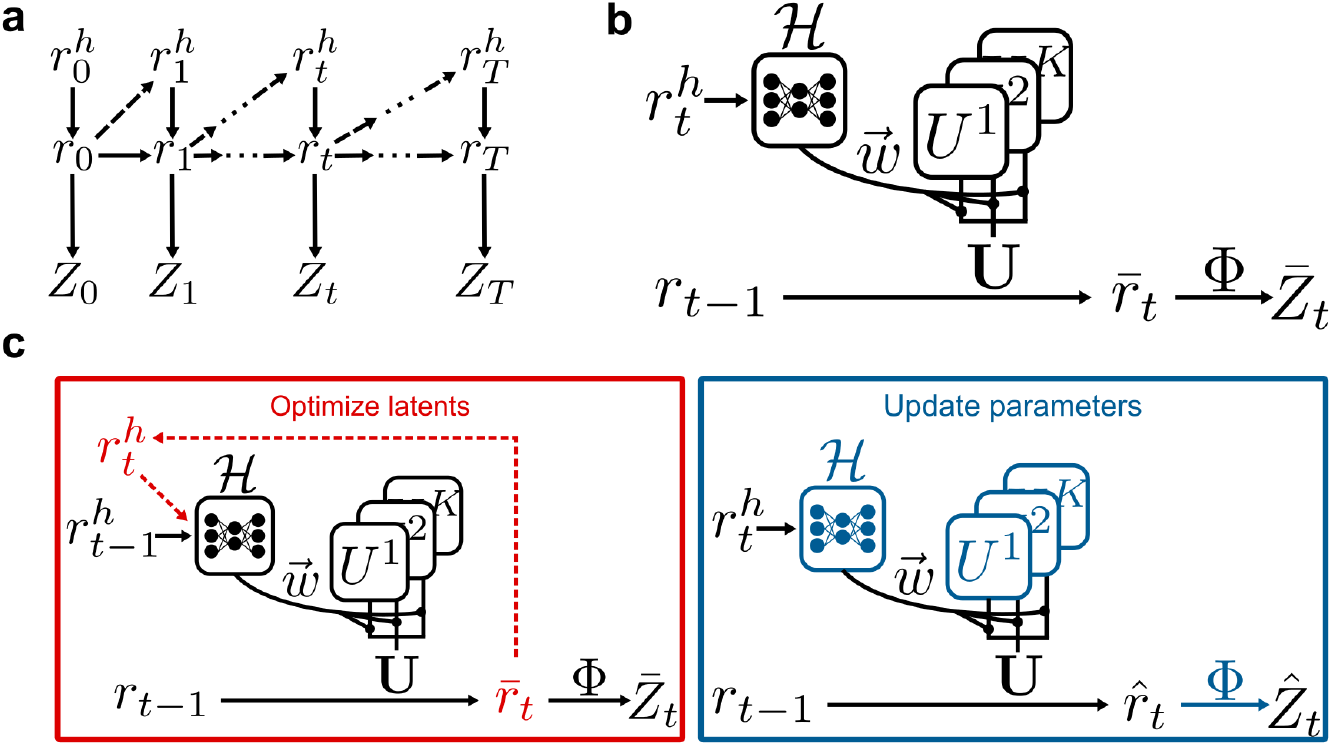
Schematics of TiDHy model and training approach. **a)** Schematic of the generative model. At every time step *t*, a higher order latent variable 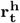 modulates the lower level latent variable **r**_**t**_. **r**_**t**_ then emits the observation **Z**_**t**_. **b)** Implementation and parameterization of TiDHy. At each timestep *t*, 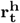 is the input to a hypernetwork ℋ, which then outputs the weight vector 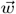. 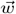 then linearly combines the set of transition matrices **U**^(**k**)^ from *k* independent latent systems. The transition matrix **U** takes the lower latent variable from **r**_**t***−***1**_ to **r**_**t**_. Finally, the observation is predicted with a observation model Φ. **c)** Schematic of training algorithm, alternating between an optimization on the latents and an update of the parameters. The free parameters learned at each stage are shown in color. At every time step *t*, an estimate of 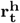 and **r** is optimized to convergence that minimizes the loss function *ℒ*_*t*_, as described in Sec. 2.2. After *T* time steps in the temporal sequence, parameters are updated with the cumulative loss and regularization (10).

The goal of TiDHy is to learn a linear combination of dynamics **U** and the latent variables **r**_**t**_, 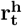, to best predict the observed data **Z**_**t**_. These dynamics could represent multiple timescales, each of which can be time-varying. After training, our method has learned a basis set of latent dynamical systems that, when combined, are able to predict observed phenomena. In other words, as shown schematically in Fig. 1b,

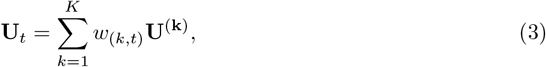

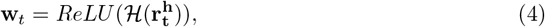

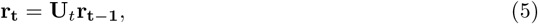

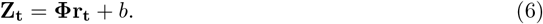

Several critical components of this modeling framework generalize and extend beyond previous methods. First, we use a flexible hypernetwork [26, 27] to model the hierarchical nonstationary dynamics. Second, while we still assume at each timestep that the dynamics are linear, this approximation is formed as a linear combination of simultaneous latent dynamics, thus capturing a hierarchy of timescales that are simultaneously active.

### 2.2 Training TiDHy

To learn a TiDHy model from spatiotemporal data, we generalize the approach from Dynamic Predictive Coding (DPC) [28], a biologically inspired normative model of predictive coding, to handle arbitrary time series data. This algorithmic procedure is conceptually similar to the expectation maximization (EM) algorithm (Fig. 1c) and is also closely related to dictionary learning and the wake-sleep algorithm [29]. In other words, each step in the algorithm first optimizes the latents and then updates the dynamic reweighing parameters, stepping until convergence. Here we describe the steps of the algorithm and the loss functions minimized in the learning procedure.

As shown schematically in Fig. 1c, in an optimization step to learn the latents, parameters *ℋ*, **U**^(*k*)^, and **Φ** are initialized and frozen. We then generate the best estimate of the latent variables **r**_**t**_ and 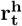, given the static parameters. For a temporal sequence of data of a predetermined length *T*, the model performs online inference of the latent variables by minimizing the sum of a dynamics predictive loss *ℒ*_dyn_ and the reconstruction loss *ℒ*_recon_, as defined below in (8) and (9). The online inference at every *t* = {0, …, T} is computed until convergence:

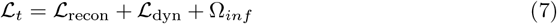

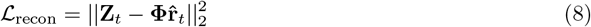

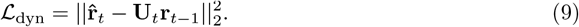

The first term ℒ_recon_ is the reconstruction loss between the optimized latent variables 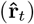 given the frozen parameters and the true observations. The second term ℒ_dyn_ compares the directly optimized lower latent variables and the estimation from the previous transition state. Finally, Ω_*inf*_ are a set of optional regularization terms for inference, such as the *l*_1_ norm of the latent variable to promote sparsity; these can be computed depending on the specifics of the data and are elaborated below.

In a second step to update the parameters ℋ, *U*^(*k*)^, and Φ (See Algorithm 1 and Fig A1 for more details), we fix the latent variables and minimize a related loss function that is summed over all time point *t* = {0, …, *T*} and combined with additional regularization terms:

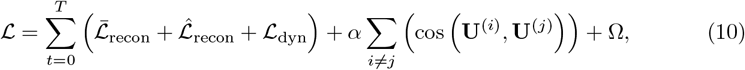

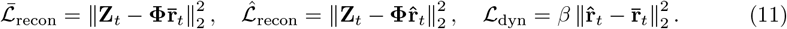

The first two terms 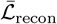 and 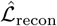 in (10) are the reconstruction losses based on the dynamic transitions 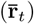, and on the optimized latent 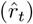. We elaborate on the use of these two different estimates of the latent variables below, and also describe how they are computed in the algorithm in Alg. 1. The third term is the predictive dynamics loss on the lower latent variables. The fourth term is a regulation term to encourage orthogonal dynamics in the solution; here we use the cosine similarity between every pair of transition matrices **U**^(*k*)^. The last term Ω represents additional optional regularization terms that can be specified during parameter updating, which we discuss below.

The two reconstruction losses are computed based on two estimates of the lower latent variable. In particular, 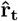 is directly optimized to make the best prediction of **Z**_**t**_ given frozen parameters, while 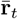 is estimated through the transition matrices 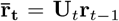, where 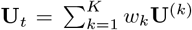. When updating the parameters *ℋ*, **U**^(**k**)^, and **Φ**, we use losses computed using both estimates of the latent variables 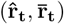 to ensure that the latent dynamics maximally reflect the ground truth observations.

We incorporated two different additional regularization terms to the overall loss function. Crucially, we added cosine regularization to the transition matrices to encourage orthogonal dynamic matrices and create an effective basis set to predict observed data. This implementation supported learning of diverse latent dynamics while also maintaining an accurate reconstruction of observables. Additional regularization can be added to the loss function during training, depending on the specifics of the data. For example, *l*_1_ norm regularization on the transition matrices can be used to encourage sparsity, allowing us to over-parameterize the dynamics with a larger *K* and still converge to a smaller model that accurately predicts the data.

The steps of the the TiDHy algorithm are summarized as pseudocode in Algorithm 1, and an implementation is available at: https://github.com/elliottabe/TiDHy. Due to the difference in scale of the terms in the loss function, we use gradient normalization [30] to stabilize training and weigh the gradients from the reconstruction loss and dynamics loss more equally. This gradient normalization occurs during parameter updates.

#### Algorithm 1

TiDHy Training

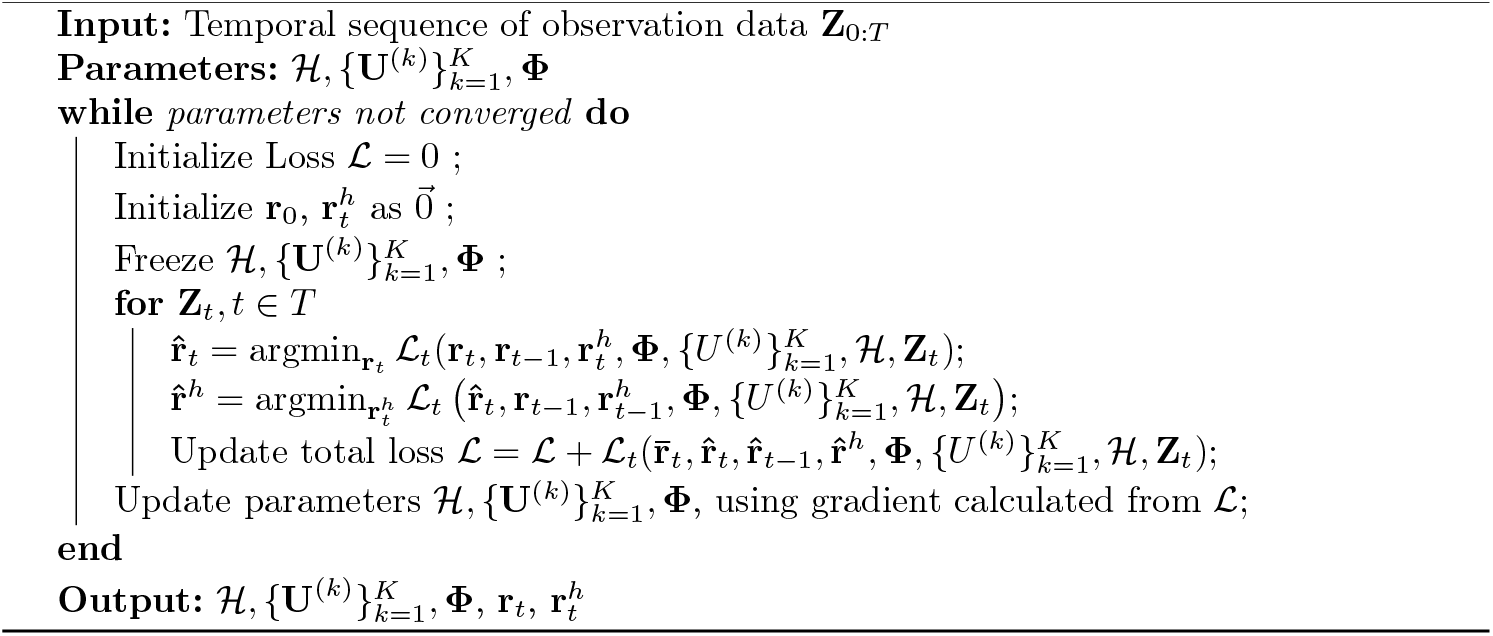

### 2.3 TiDHy demixes multiple independent latent dynamics

To assess TiDHy’s ability to capture latent dynamical systems, we first designed a synthetic dataset to represent complex, noisy signals with latent dynamics. To generate a dataset with distinct latent timescales, we simulated 3 independent 2-dimensional SLDSs (*p* = 1, 2, 3) with different eigenvalues for each set of latent dynamics, where each system has both a fast and slow dynamic matrices (*k* = 1, 2), for a total of 6 latent dynamical systems of which any 3 can be active at any time. The generated emissions were then partially superimposed and randomly projected into a higher-dimensional space to mimic experimental data (Fig 2a). Each SLDS system follows the standard formulation:

**Fig. 2.**
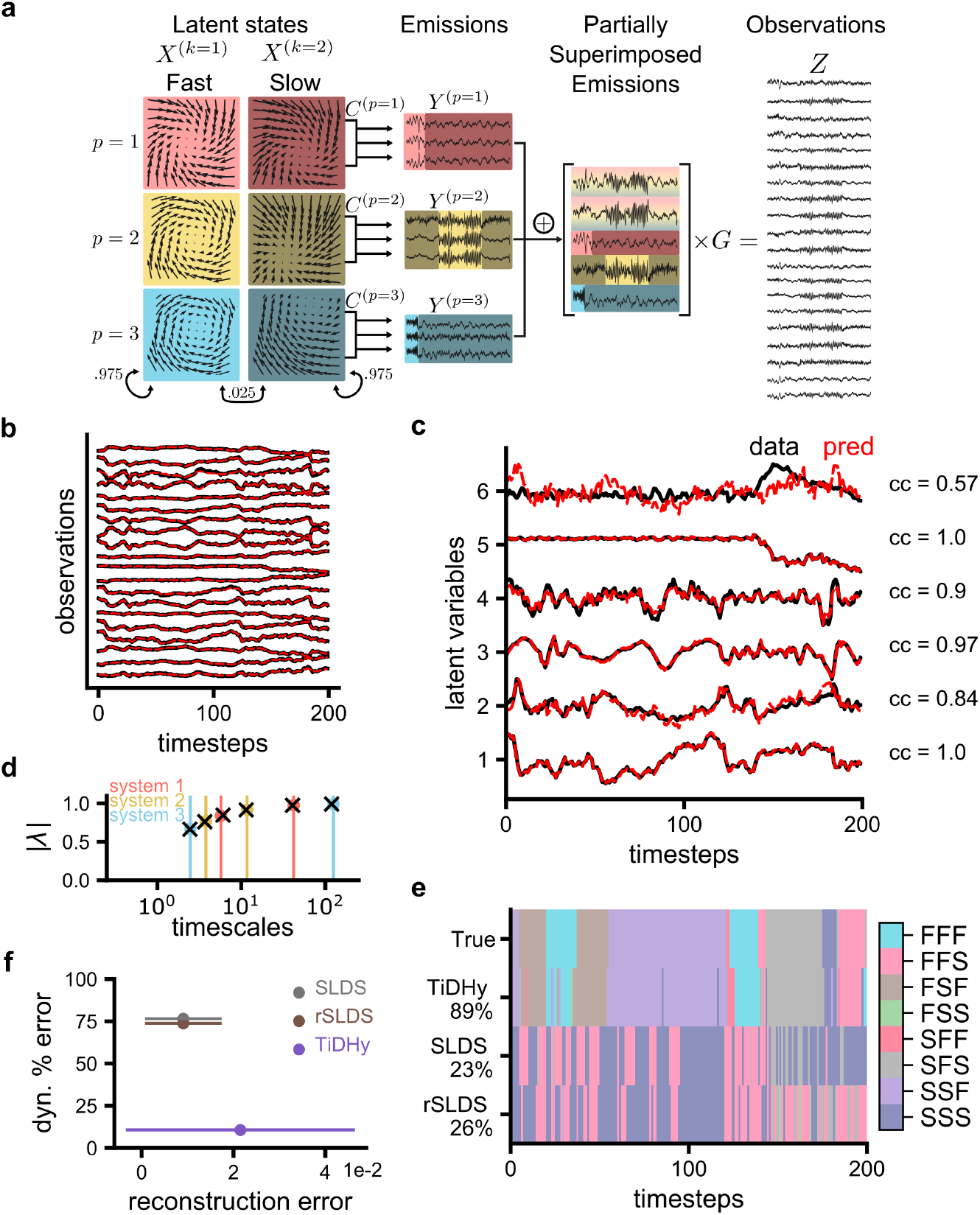
TiDHy demixes multple simutaneous SLDSs in synthetic dataset. **a)** Schematic of how the synthetic dataset was generated. Three independent SLDSs were used to generate emissions coming from either a set of slow or fast dynamics. The emissions from each system were then partially superimposed and projected into a higher dimensional space. **b)** Comparison of ground truth data (**Z**_**t**_, black) and predicted observations (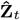, red). The mean error between the ground truth and prediction observations was 0.016±0.015. **c)** TiDHy learns latent spaces that closely match the ground truth. Since the latent space is degenerate up to an arbitrary rotation, we used CCA to compare the components of the ground truth (**X**_*t*_, black) and the lower-level latent variable (**r**_**t**_, red). **d)** The timescales captured by the lower-latent variable **r**_**t**_ match those of the ground truth. The timescale is computed by fitting an AR model to the lower-latent variable **r**_**t**_ and calculating the corresponding eigenvalues. The y-axis represents the magnitude of the eigenvalues *λ*. The vertical lines represent the true eigenvalue timescale of each system and colored dots represent the magnitude of the true eigenvalues. Black X’s represents the closest timescales captured by TiDHy. **e)** TiDHy accurately predicts which combination of simultaneous systems is active, out performing a single large SLDS model. Each color represents a unique combination of dynamics (e.g., FSF = fast, slow, fast). The accuracy of the KNN classification is shown on the y-axis. **f)** A comparison of the dynamic identification error against the reconstruction error. While TiDHy does not reconstruct the observations as well as the rSLDS and SLDS model, it captures the true simultaneous latent dynamics with higher accuracy. Error bars are standard deviation of the reconstruction error.

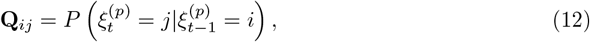

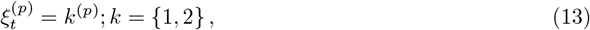

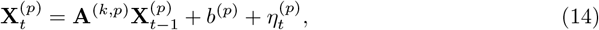

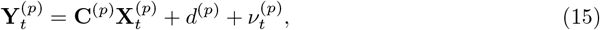

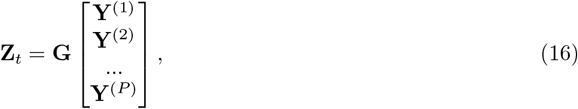

where the superscripts (*k, p*) represent the indices for the dynamics matrix and system number, respectively. Each system switches between a fast (**F**, *k* = 1) and a slow (**S**, *k* = 2) mode based on a hidden Markov model (HMM) with probability **Q**_*ij*_ = 0.025 in its latent dynamics 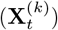 (See Methods). Thus, with three systems of two timescales each, there are eight combinations of unique simultaneous timescales active at any given time (e.g., FFS, FSF, SFF, etc.). The latent dynamics 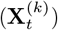 of each system are two-dimensional rotations that evolve according to dynamic matrices **A**^(*k*,*p*)^, with an emissions model **C**^(*p*)^. The emissions of each independent SLDS are partially superimposed by summing the emissions and randomly projecting them with **G** into a higher-dimensional space (**Z**_**t**_) (Fig 2a).

TiDHy is trained to learn the latent dynamics by predicting the mixed observation data using the algorithm described in Section 2.2. After training, we first verified that TiDHy correctly predicts the input data (Fig 2b). On average, the Mean Squared Error is 0.016±.0.015 a.u. More importantly, we next verified that TiDHy learned the correct latent dynamics by correlating the learned latent variables (**r**_**t**_) with the ground-truth latent variables (**X**_*t*_). Since we do not constrain the structure of **r**_**t**_, we can learn any rotation of the latent space that produces the same prediction of the observations. To account for this, our evaluation used canonical correlation analysis (CCA) to align the learned and ground-truth latent variables (Fig 2c). We find that lower-level latent variables **r**_**t**_ are highly correlated with ground truth latent dynamics.

To examine whether TiDHy could capture the ground truth timescales, we also analyzed the eigenvalue structure of the learned dynamical systems. Because the model learns onetime step predictions, the transition matrix **U**_*t*_ can have unstable modes that do not grow exponentially and thus do not represent the full dynamics over an entire sequence. Thus, to verify that the model captures the correct eigenvalues over the data sequence, we first fit an auto-regressive (AR) model on the latent variables **r**_**t**_. In the case of the SLDSs, we have ground truth knowledge about when the systems are in either a fast or slow state. By fitting the AR model on data when each state is active and then computing the timescales (see Methods), we show the eigenvalues of the AR model closely represent the ground truth timescales of simulated latent dynamics (Fig 2d).

Finally, to verify that each of the simultaneous timescales within the dataset is correctly captured, we use a K-nearest neighbor (KNN) approach to classify the unique combination of timescales active based on the higher-order latent variable 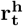 (Fig 2e). To benchmark TiDHy, we compare this classification accuracy against two leading alternative approaches to spatiotemporal demixing, namely a single large SLDS and a recurrent SLDS (rSLDS) that switches among eight transition matrices that can be switched between (i.e., the rSLDS can learn up to eight unique combinations of timescales) [10]. TiDHy consistently outperforms the traditional SLDS method, which parameterizes multiple systems as a single large dynamical system (Fig 2e,f). Over the entire simulated dataset, TiDHy can accurately classify the unique set of timescales at each time step 89% of the time in comparison to 23% and 26% for SLDS and rSLDS respectively (Fig 2e). Crucially, the SLDS can reproduce the observations but is unable to accurately capture the true simultaneous latent dynamics (Fig 2f).

One of the benefits of a generative model such as TiDHy is the ability to make accurate predictions of new data. In the training procedure of TiDHy, it is necessary to specify a temporal window of length *T*; however, by creating overlapping windows, our method can learn dynamics across longer timescales outside of a single window length. This is particularly beneficial when experimental data is continuous and not organized in trial formats. We show that even when trained on a limited window size (trained with *T* = 200), TiDHy can still accurately predict the observations (Fig 3a) and learn the latent variables **r**_**t**_ (Fig 3b). TiDHy also correctly classifies the unique simultaneous dynamics (Fig 3c,d). Using a shorter window length *T* than originally trained results in only a small degradation of the KNN classification accuracy (Fig 3d).

**Fig. 3.**
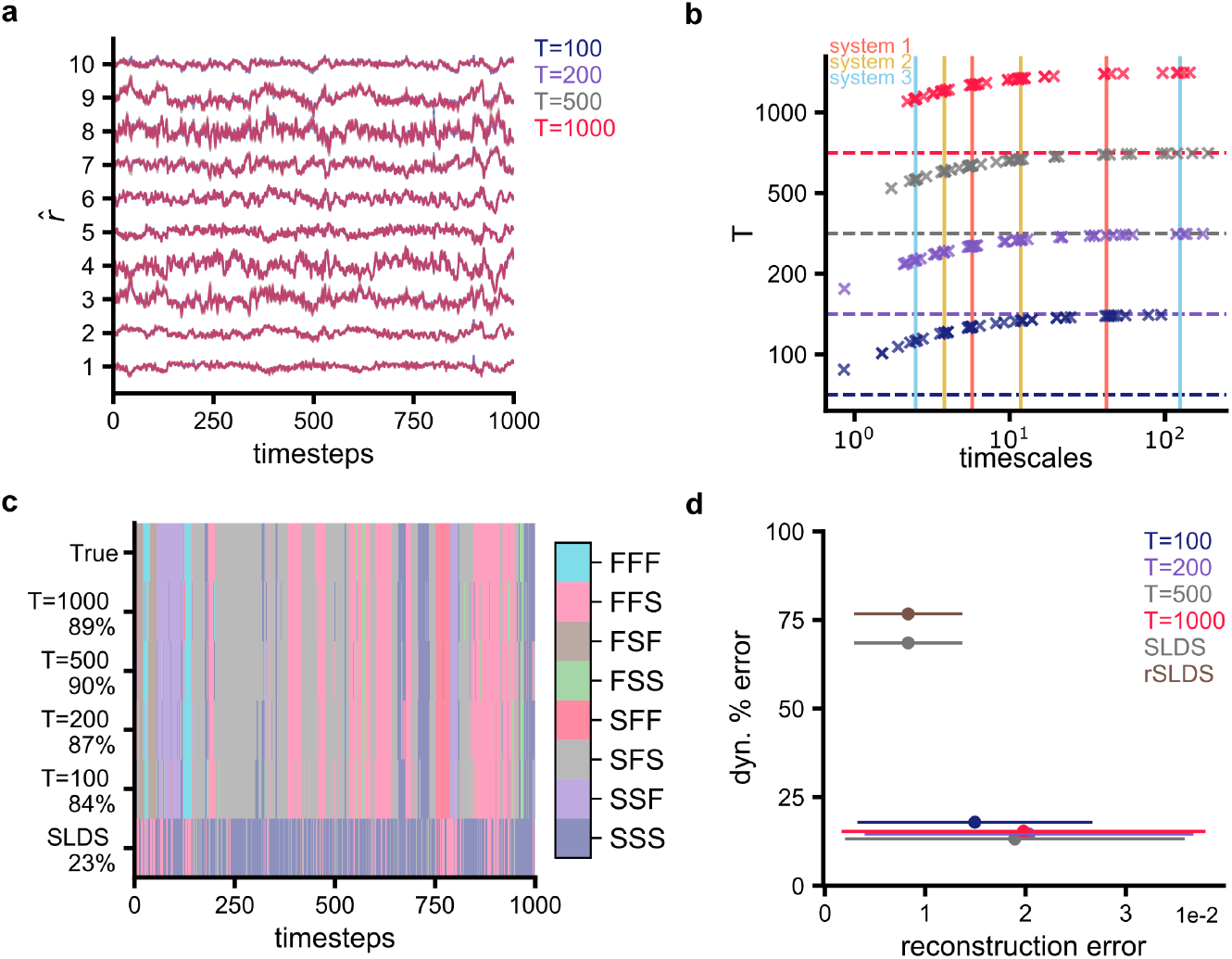
TiDHy captures stable simultaneous timescales. **a)** Comparison of latent variable **r**_**t**_ trained using different window lengths *T* = *{*100, 200, 500, 1000*}*. **b)** Comparison of timescales calculated from AR model fit on the latent variables **r**_**t**_. Each row represents different temporal window lengths on which TiDHy was tested. Within a row, the values represent the magnitude of the eigenvalues like Fig. 2d. **c)** KNN classification of simultaneous unique combination of timescales tested on 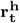 from TiDHy models using different sequence lengths. The accuracy of the classification is shown on the y-axis. **d)** TiDHy model trained with *T* = 200 and tested with other window lengths. A comparison of the dynamics identification error against the reconstruction error. Even when extending the window length, TiDHY can reconstruct the observations with the same accuracy while accurately identifying the true latent dynamics. Results are from N=8 random seeds. Error bars are standard deviations of the reconstruction error.

### 2.4 Learning slow timescales in simulated locomotion

To test TiDHy on a behavior dataset with multiple timescales where the ground truth dynamics are still well known, we used a simulated locomotion dataset [31, 32] (Fig 3a). In this simulated behavior, we show that TiDHy can simultaneously identify the fast dynamics of the kinematics and slow transitions as the robots walk across different terrains.

The dataset contains movement kinematics of two quadruped robots, ANYmal B and ANYmal C, simulated while walking on diverse terrains in NVIDIA’s Isaac Gym [33] physics simulator. The environment has five different terrain types (namely: flat, incline stairs, decline stairs, incline slope, and decline slope) that are procedurally generated. The arena of the dataset is simulated on a 12×12 grid of 1 meter squares that are randomly sampled from the five terrain types. In addition to different terrain types, each square is parameterized by roughness, slope, and a discrete value for level of difficulty. In the context of demixing multiscale data, each robot’s unique morphology represents a global timescale, the terrain is a medium timescale, and the rhythmic locomotor kinematics is a fast timescale. The dataset contains the kinematics of joint angle and velocities recorded over 5182 walking trajectories. These kinematics are similar the type of data that is most commonly typically experimentally accessible with keypoint tracking, so they are used as observations to train TiDHy.

We evaluated TiDHy on metrics similar to those we used with the synthetic dataset in Section 2.3 and showed that it is able to learn all three relevant timescales in this locomotor dataset. We first verified that the model could accurately reconstruct the dynamics of the observation data (Fig 4b). We then performed similar analyses on the latent variable **r**_**t**_ by fitting an AR model and calculating the timescale of the dynamics (Fig 4c).

**Fig. 4.**
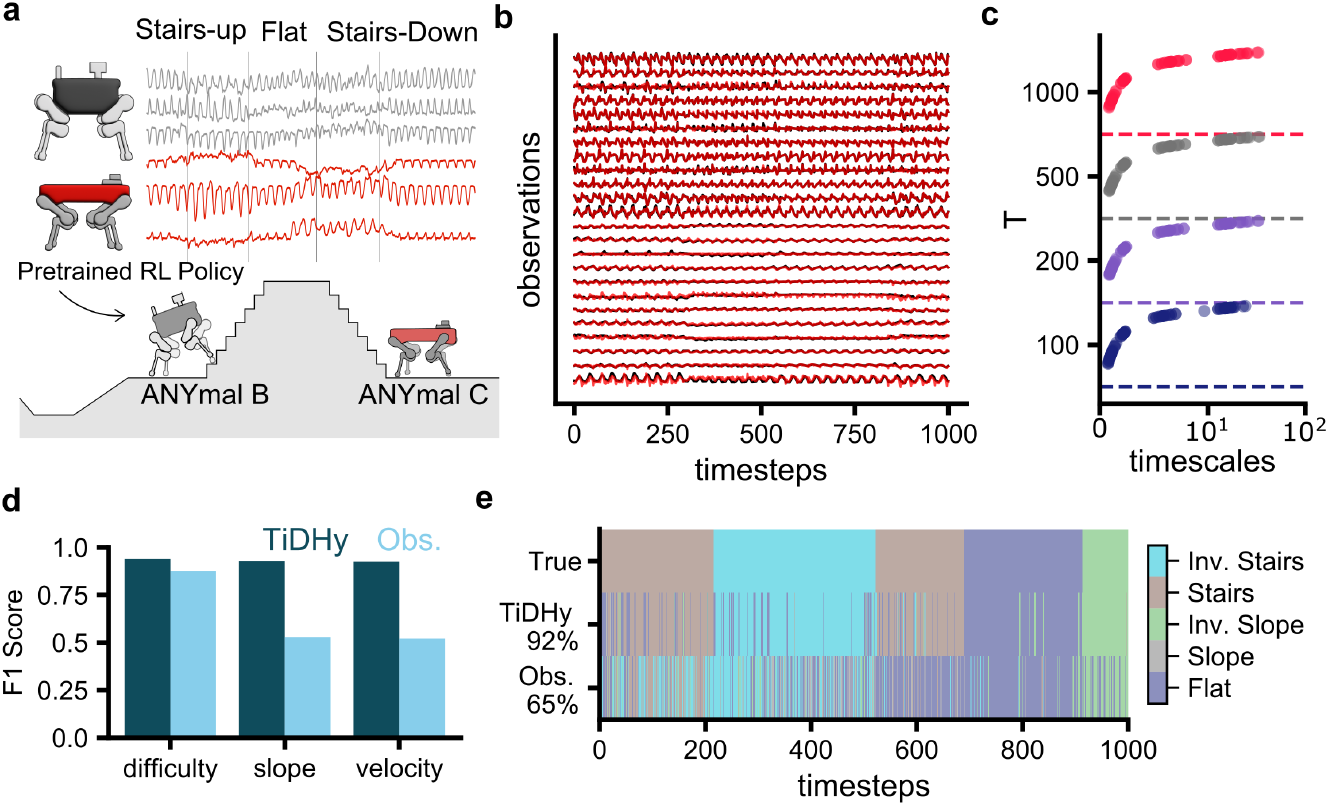
TiDHy learns the simulaneous dynamics of simulated locomotion. **a)** Schematic of the simulated locomotion dataset. Two robots with different morphology are simulated walking on different terrains in NVIDA IssacGym. Schematic modified from [31]. **b)** Example of input observations used to train TiDHy. Black traces are the real kinematic data from the simulations. Red traces are predictions from TiDHy. **c)** Timescales of the latent state **r**_**t**_. TiDHy trained on T=200 and tested on additional time windows. The timescales are computed from an AR model fit to the latent variable **r**_**t**_. **d)** Using the higher order latent variable 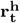, TiDHy can correctly classify the within-task variables like the difficulty of the terrain, slope, and command velocity compared to using the observation data directly. **e)** KNN classification of the type of terrain the agent was walking on across time. TiDHy correctly identifies the terrain against a comparison by using the observation data directly. Percentages represent the accuracy of the classification.

Similar to the previous section, we first trained on temporal sequence lengths of *T* = 200 timepoints and tested on temporal windows of size *T* = 100, 200, 500, 1000. These timescales remain consistent across testing of different max sequence lengths. From this analysis we find three main timescales of the data, that represent the faster joint dynamics, the changes in the terrain and length of the simulations as a proxy for body morphology.

Next, we wanted to explore the type of structures the higher-order latent variables 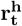 captured. In this analysis, we could identify the task level details using the demixed latents learned from kinematic observables, such as the value of the slope at each frame, the category of difficulty, as well as the robot’s command velocity (Fig 4d). In comparison, direct observations of the joint dynamics was not able to capture these variables.

To verify that the variables we extracted were a latent part of the system, we also compared our model’s ability to classify the task properties with the same classification computed directly from the joint angles and velocities (Fig 4d,e). Fig. 4e shows a single robot as it traverses different terrains overtime. Using the higher-order latent variable 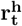, TiDHy correctly identified the terrain type the robots were walking on using the KNN classification (Fig 4e). TiDHy accurately predicts the terrain with 92% accuracy, while in comparison to using the direct joint dynamics achieves 65% accuracy. These results demonstrate TiDHy’s ability to capture real latent information that is interpretable as meaningful quantities of the system.

### 2.5 TiDHy captures latent dynamics in social behavior

Next, we applied our method to explore an open source mouse social behavior dataset, CalMS21 [34]. This social behavior dataset was particularly interesting as a test case because there is no trial structure (i.e., each clip of the mice is variable in length). Additionally, the dataset contains data from two systems (i.e., each mouse) that can act independently or may be coupled together. This behavior tracking dataset has positions of seven keypoints on each mouse: the nose, the left and right ears, left and right torso, and the base of the tail. The published dataset also contains annotations from four experts for three distinct social behaviors (‘attack,’ ‘investigation,’ ‘mount’) and an ‘other’ category. We trained TiDHy by directly fitting ~300 minutes of keypoint tracking data, during which time two mice were interacting, and demonstrate TiDHy’s flexibility in capturing and demixing the latent dynamics in natural behaviors.

We first confirm that TiDHy fits the data accurately (Fig 5b). TiDHy was able to accurately predict the keypoint data of the mice with an average error of 0.04 *±* 0.03 pixels. Next, we perform a timescale analysis on the latent variable **r**_**t**_ by fitting an AR model, thus demonstrating that there are distinct timescales present during social interactions (Fig 5c). The model was trained with a temporal window *T* = 200, and then we extended or contracted the temporal window to investigate how stable the learned temporal dynamics were. By clustering of the learned timescales, we estimate that there are three major timescales in the social behavior, on the order of tens of seconds, hundreds of seconds, and the last on the order of tens of minutes. We speculate the first timescales represent individual movements of the mice, while the timescale in the tens of seconds represents the behavioral syllables. Finally, the last timescale likely represents the length of the clips recorded in the dataset.

**Fig. 5.**
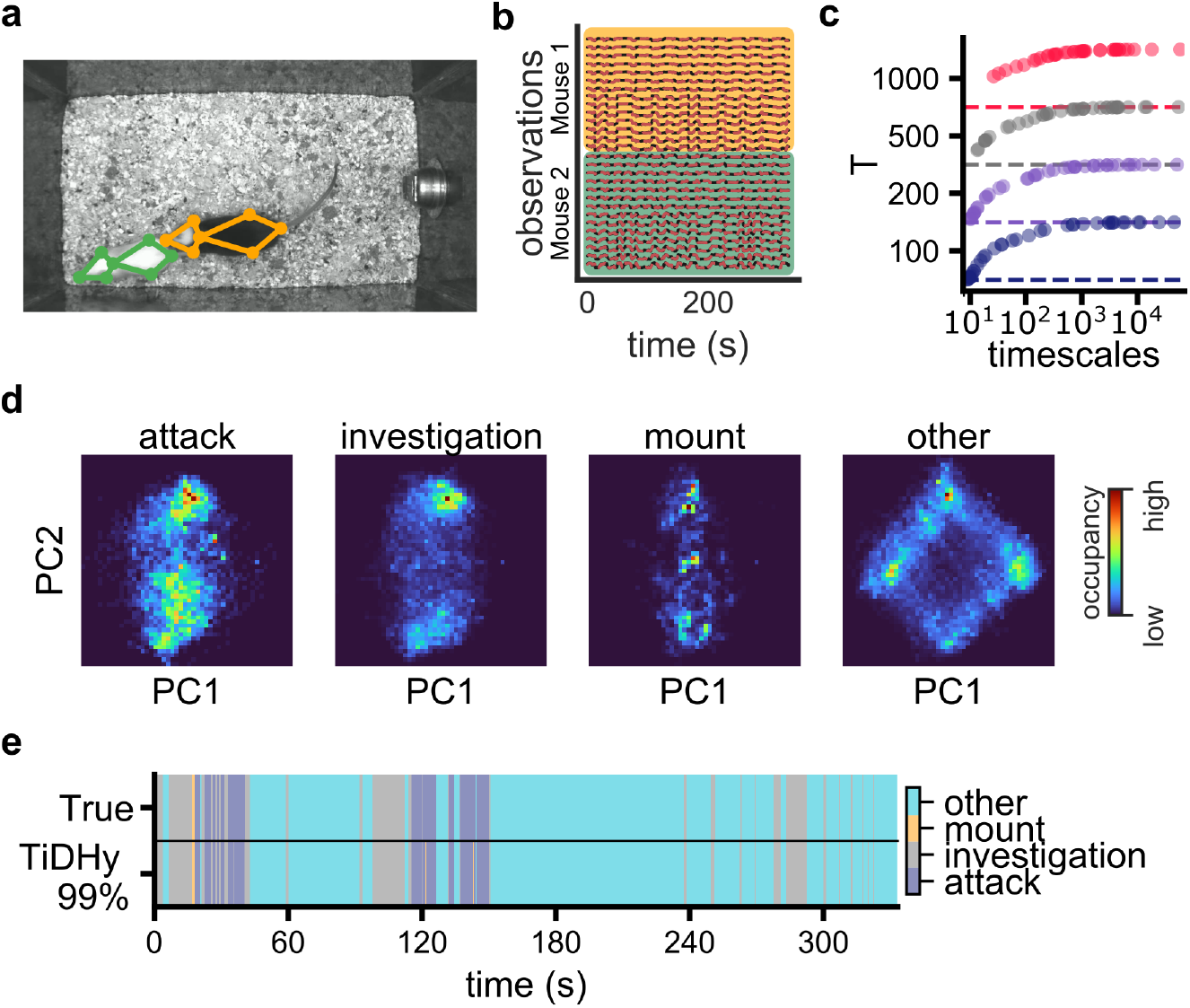
TiDHy captures the behavioral dynamics of interacting mice. **a)** Example frame and key point tracking from the CalMS21 dataset. **b)** Example of the observations used to train TiDHy. The black traces are the ground truth key point data and the red are predictions from the trained model. The distinction of each mouse is only used for visualization and not used as input into the model. **c)** To gain an intuition of the timescales of the behavior, we fit an AR model on the latent variables **r**_**t**_ and computed the timescales. The model was trained with a temporal window of *T* = 200, and tested on other window sizes. We find stable timescale information independent of window size. **d)** PCA of the latent variable **r**_**t**_ split by the expert annotations. We find the first PCs accounts for when the mice are interacting. Colormap is a 2d histogram representing the proportion of time spent. **e)** KNN classification of the higher-order latent variable 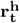 accurately identifies the expert annotated behaviors.

Next, we show that TiDHy creates an interpretable latent space, as visualized by the top two principal components (PCs) of **r**_*t*_ as a 2D occupancy histogram. In visualizing the data using expert behavioral annotations, we observe that the learned representations of these labeled social behaviors lie on a specific hyperplane in the latent space (Fig 5d). Based on this analysis, TiDHy separates the representations of individual mice when they are in the ‘other’ state (typically when they are ignoring each other). Using this approach, future work can explore different clustering methods to partition the latent space and redefine these course behavioral labels. Finally, TiDHy can recapitulate the ground truth behavior labels using KNN regression on the learned latent space of 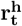 with high accuracy (99%, Fig 5e).

## 3 Discussion

Motivated by the need to model an increasing complexity in spatiotemporal recording of natural behaviors, we develop TiDHy as a method for demixing simultaneous, multiscale latent systems. TiDHy learns to demix data by learning a hierarchical generative model, where linear combinations of dynamics are modulated from timestep to timestep. We use a hypernetwork to learn a set of latent dynamics, so that they are capable of spanning orders of magnitude different timescales. By examining its performance on several synthetic and experimental datasets, we demonstrate TiDHy’s ability to learn multiscale latent dynamics in a range of contexts and data types. Importantly, we evaluate TiDHy not only on reconstruction of observed data alone, but also on its ability to learn the structure of the true latent dynamics. Further, by creating multiple estimates of the latent representations, TiDHy converges to stable latent dynamics that can be extended to continuous non-trial structured data by changing the size of the temporal window. While we primarily developed TiDHy for investigating recordings of kinematics during natural behaviors, we envision our method as a general-purpose tool for demixing timescales from timeseries data. Future extensions of the model can incorporate more complex architectures such as variational autoencoders [35, 36], representational losses to constrain the structure of latent spaces [31, 37], or joint embedding of neural and behavioral data [38, 39].

### 3.1 Applications to modeling natural behavior

From a swift escape from a predator to hours foraging for food, natural animal behaviors are characterized by latent dynamics that span multiple spatial and temporal scales. For example, when a mouse seeks food, its decision to search in a particular area is influenced by slow dynamics, such as satiety or whether they visited the area recently, and also fast sensory cues, like concentration of odors, presence of predators, and the terrain of the environment. Both timescales jointly contribute to explain total variation in behavior. Historically, due to the limitations of technology, systems neuroscience has focused on head-fixed, trial-based experiments. However, trial-based data only capture a small portion of an animal’s behavioral repertoire. With recent developments in computer vision [40–42] and neural recordings [43], datasets that contain increasingly naturalistic behavioral and neural recordings are becoming possible [7, 44–48]. We developed TiDHy to meet the challenge of parsing and modeling such multiscale neuroethological data.

One of the key characteristics of natural behavior is the lack of a strict trial structure. This presents a challenge for many methods [9, 10, 49] as it is common practice to process data with a fixed window of time. While TiDHy also uses a fixed temporal window during training, when applied to new data, we show in Section 2.3 how the temporal window of the input data can be expanded or contracted to the specifics of the dataset and still yield consistent dynamics and timescales. This feature has several benefits, including reduced computational resources for training and flexibility in applying it to many different types of timeseries data. For example, in natural behaviors, multiple animals can interact and TiDHy can disentangle and interpret how the behavior of each animal evolves over time.

### 3.2 Predictions as a learning signal

Both biological and artificial systems can use predictive processing as an efficient way to generate a learning signal. The ability to predict the future is a fundamental computation that is critical to the survival of complex biological systems. For instance, predictions are essential to construct spatial maps [50], represent sensory information [28, 51, 52], and motor control [53]. These representations are crucial for ensuring that an animal can respond flexibly and efficiently to changes in the environment. Similarly, training predictive models in artificial systems have been fruitful in building semantic representations of videos [54], speech [55, 56], and reinforcement learning [57, 58]. In training TiDHy, we leverage this efficient learning approach to construct a predictive loss function to learn latent dynamics from arbitrary timeseries data.

Mapping partial observations to latent variables is an inverse problem with inherent ambiguity. In other words, multiple latent dynamics can yield identical predictions, making it difficult to discern the true underlying dynamics solely based on observed data. To address this issue, we constrain the possible solutions by imposing a linear transition between time points while balancing predictions from the optimized latent variables 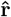 and the dynamics 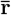. This constraint helps to reduce the solution space, guiding the model towards more plausible and interpretable representations of the latent dynamics. By enforcing linear transitions with modulation from the hypernetwork, we simplify the relationships between consecutive time points, facilitating the convergence to stable dynamics. Additionally, balancing the predictions from 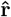 and 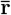 (Eq. 10) ensures that the model considers both the optimized latent variables and the overall dynamics, leading to a more robust and reliable learning process.

### 3.3 Limitations and assumptions of TiDHy

To reduce the computational time needed to train TiDHy, we have defined a temporal window to compute the latent dynamics. While TiDHy can learn timescales beyond a single temporal window, the model still requires a specified window length during training that is long enough to capture some of the timescales within the data. Additionally, the core of the model represents simultaneous dynamics as linear combinations and transitions between time points, relegating the nonlinearity to the hypernetwork. As a result, there is a tradeoff between the expressibility of the latent space and the interpretability of the representations. In the scope of this work, we have limited ourselves to favor interpretability with linear latent spaces, but an extension of TiDHy can also use nonlinear representations. Additionally, while TiDHy can learn multiple simultaneous dynamics, we restricted our analysis to the continuous domain. There are other methods [59, 60] that can be added to the analysis to discretize the latent representations for system identification after training. Finally, when training and evaluating TiDHy, the dimensionality of the observed data needs to remain constant. A potential solution to missing observations (e.g., changing number of animals in view of tracking cameras) is to retrain the observation model while keeping dynamics and hypernetwork fixed.

Recently, a method was introduced with the similar goal of decomposing linear dynamical systems (dLDS, [14, 15]); TiDHy is related to dLDS, but they have different conceptual motivations and implementations. Decomposing LDS uses sparse dictionary learning to learn a set of coefficients with a linear combination of dynamics. However, the motivations for dLDS and TiDHy differ fundamentally. dLDS was specifically developed as a descriptive data model, whereas TiDHy is designed as a hierarchical generative model. We specifically chose to model observed data as a hierarchical generative model to enable future work with real-time analysis, prediction of unseen data, and control. Furthermore, there are two main implementation differences between TiDHy and dLDS. First, we use an additional higher-order latent variable 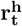 to learn a nonlinear mapping via a hypernetwork to modulate the linear combination of dynamics. Second, our optimization procedure uses a predictive loss that does online correction and, as a result, can be used on new unseen data. These two factors enable TiDHy to learn not only non-stationary dynamical systems but multiple simultaneous ones.

### 3.4 Extensions of TiDHy

There are several direct extensions of the model that can be made to improve interpretability and learned representations. As datasets grow in size and complexity to incorporate simultaneous neural activity during natural behaviors, methods to understand how neural and behavioral data jointly interact are needed. Some methods have recently attempted to tackle this [38, 39]. However, these models utilize complex neural networks to compress the data into a latent space with highly nonlinear mappings. Instead, for better interpretability, TiDHy could be adapted to create a joint embedding space where neural activity informs behavioral changes. A potential solution is to use neural activity as an input to the hypernetwork modulating the latent dynamics of behavior. Alternatively, the inputs could switch where the inputs to the hypernetwork are behavioral measures that modulate the latent dynamics of the neural activity.

Currently, our implementation only uses the temporal dynamics loss to learn the higher-order latent variable 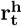. To learn more structured latent spaces, it is possible to include additional constraints on 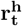, such as representation auxiliary losses [31, 37]. In addition, other than encouraging orthogonality of the latent dynamics with cosine similarity, the learned dynamics are not explicitly constrained; as a result, they can be modified for specific datasets. For example, low-rank tensor operations have been shown to be an effective way to make interpretable and expressive representations [61–65]. In this work, we restricted the model to utilize linear decoding to predict the observed data. A modification to allow nonlinear transformation of the observed data would implement a variational autoencoder (VAE) [66] architecture.

## Acknowledgments

We thank Anna Bowen for discussions, feedback, and for coining the acronym TiDHy. We thank Linxing Preston Jiang for discussions and code on DPC. Additionally, we thank the members of the Brunton lab for their feedback on drafts of the manuscript and discussions throughout the project.

This work has been funded through a Swartz Foundation Fellowship in Theoretical Neuroscience to ETTA; AFOSR award FA9550-19-1-0386, NIH U01NS136507, and the Richard & Joan Komen University Chair to BWB.

## Author Contributions

E.T.T.A. conceived the project, designed the experiments, implemented the algorithm, and performed analyses. E.T.T.A. and B.W.B wrote the manuscript.

B.W.B supervised the project.

## 4 Data and Code Availability

Code to train TiDHy and generate SLDS data can be found at https://github.com/elliottabe/TiDHy. The data for the simulated locomotion behavior is from [31] and can be found https://github.com/nerdslab/bams simulated quadrupeds. The CalMS21 dataset [34] are available at https://data.caltech.edu/records/s0vdx-0k302.

## 5 Methods

### 5.1 Additional details on TiDHy model

The hierarchical generative model follows the procedure presented in [28]. The set of hyperparameters for general training can be found in Table 1. The form of Ω for regularization was kept to a minimum in all fitting procedures. To help disenangle the higher-order latent variables 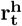, we implemented a cosine regularization on the input weights to the hypernetwork. This ensured the encouraged the representations across time for 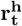 were approximately orthogonal.

**Table 1.** Table of Model Hyperparameters.

| | $\mathcal{H}$ | $U$ | $\Phi$ | $\mathbf{r}_t$ | $\mathbf{r}_t^h$ |
| --- | --- | --- | --- | --- | --- |
| Learning Rate | 1e-3 | 1e-2 | 1e-2 | 2e-1 | 2e-1 |
| Dimensionality | 50 | 15 | (10 x input_dim) | 10 | 5 |
| Optimizer | AdamW | AdamW | AdamW | SGD | SGD |
| L1 Regularization | 0 | 0 | 0 | 0 | 0 |
| Cosine Regularization | 0 | 0 | 0 | 0 | 5 |
| L2 Regularization | 1e-5 | 1e-5 | 1e-5 | 0 | 0 |

After training, we used Scikit-learn’s K-nearest neighbor classification to decode the relevant information from the latent states 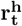 and **r**_**t**_.

The training was done on a single NVIDIA RTX A6000 GPU.

### 5.2 Multiple SLDS Dataset

To generate the independent SLDS systems, we utilized the SSM python package [10]. The latent dynamics are two-dimensional rotations with switching dynamics between a fast and slow mode. The switching probabilities were governed by a hidden Markov model where the probability of switching for each system was *Q*_*ij*_ = 0.025 and staying in the same state was *Q*_*ii*_ = 0.975. The eigenvalues of the rotation matrix of the latent dynamics were empirically chosen to provide non-overlapping timescales. We simulated a train, validation, and test dataset with 200,000 and 50,000 time steps. Each system was simulated independently, and the emissions were concatenated, partially superimposed, and then randomly projected into higher-dimensional space. To preprocess the data, we normalized by the max of the absolute value. The timescales of the latent systems were calculated by first fitting a one-step auto-regressive model (AR) to the latent variables **r**_**t**_. Next, we computed the timescale as 1/|*Re*(*ln*(*λ*)) |, where *λ* are the eigenvalues of the AR model.

The additional hyperparameters are found in Table 2.

**Table 2.** Training parameters.

|  |  |
| --- | --- |
| Nepochs | 1,000 |
| Total Time Train | 200,000 |
| Total Time Val/Test | 50,000 |
| Temporal Sequence Length | 200 |
| $\alpha_{cos}$ | 5 |
| $\alpha_{gradnorm}$ | .5 |
| Overlap Factor Partial Sup. | 2 |
| Observation Dimensions | 20 |
| Overlap Window Fraction | .9 |
| $\beta$ | 2 |

**Table 3.** Training parameters.

|  |  |
| --- | --- |
| Nepochs | 1,000 |
| Total Data Train | 1000*9000 |
| Total Data Val/Test | 50*9000 |
| Temporal Sequence Length | 200 |
| $\alpha_{cos}$ | 5 |
| $\alpha_{gradnorm}$ | .5 |
| Overlap Factor Partial Sup. | 2 |
| Observation Dimensions | 20 |
| Overlap Window Fraction | .5 |
| $\beta$ | 2 |

**Table 4.**
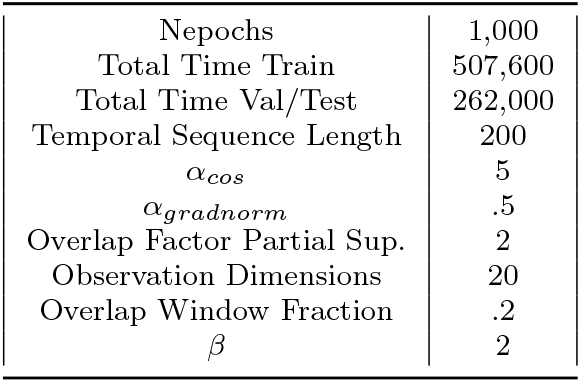
Training parameters.

### 5.3 Simuluated Locomotion Dataset

We used an open-source dataset from [31] that utilized the Nvidia Issac Sim physics engine [33]. 24 proprioceptive features, including joint position and velocities, were used, and no preprocessing was necessary for model fitting. See https://github.com/nerdslab/bams simulated quadrupeds for additional details.

To decode task variables from TiDHy, we used a KNN classification for each timepoint at during the simulations.

### 5.4 CalMS21 Datset

The details of the CalMS21 dataset can be found in [34] and https://data.caltech.edu/records/s0vdx-0k302. Only minor preprocessing was needed to train TiDHy. The data was split according to the original paper. First, we selected trials with at least *T* = 200 time points to train TiDHy. Then, the data was normalized to the key point trajectories to be between 0-1 and concatenated them into a single two-dimensional array. A total of 28 features were used as the observations to train TiDHy, representing the xy key point positions of both mice.

## Appendix A Supplementary figures

**Fig. A1.**
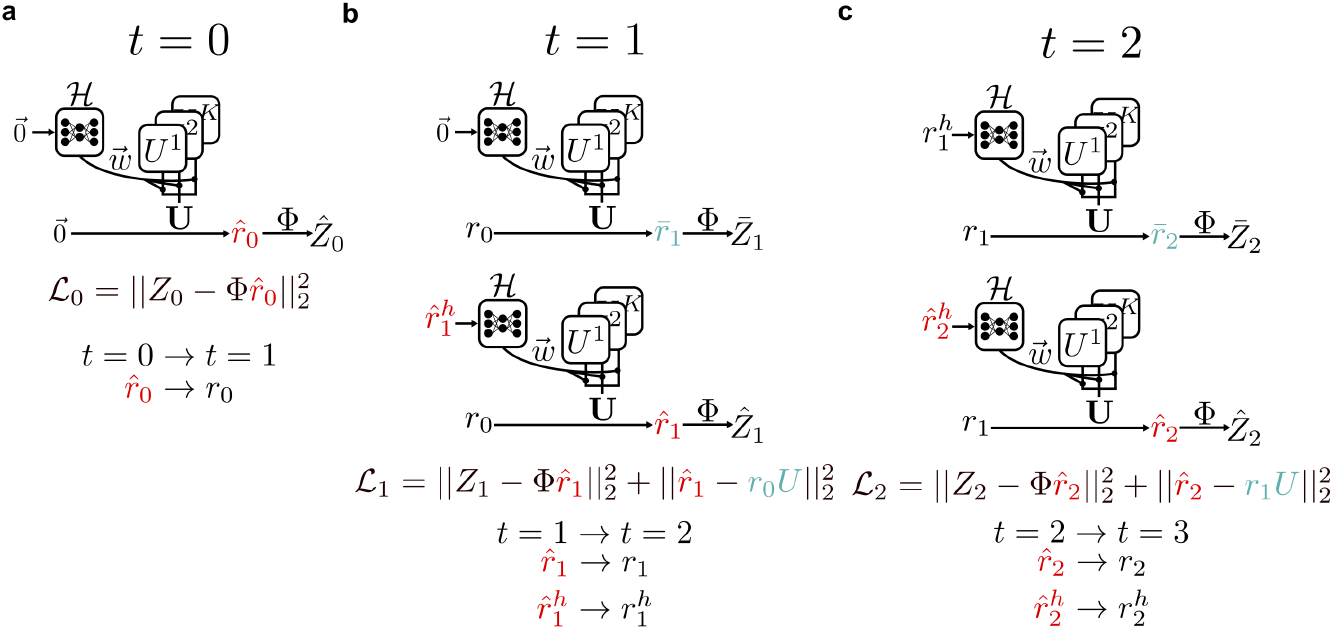
Three example time points of training TiDHy. **a)** On initialization 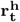 is set to 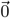 and 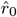 is optimized (red) to give the best estimate of the observation *Z*_0_. In the transition to *t* = 1 the optimized 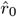 is used as the *r*_*t*−1_ for the next time point t = 2. **b)** At t ≥ 1 two versions of lower latent variables are simultaneously estimated. 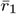 is computed using the transition matrix *U* (blue) and with the previous 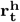. 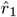 and 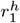 are simultaneously optimized to make the best prediction of *Z*_*1*_ given the current 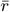. In the transition between time points the optimized latent 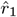 becomes the *r*_*1*_ used to estimate 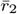. **c)** Same procedure as in b. At the end of the temporal sequence of length T, the loss values are collected and used to update the parameters *ℋ*,*U*, and Φ.

